# Global adaptation confounds the search for local adaptation

**DOI:** 10.1101/742247

**Authors:** Tom R. Booker, Sam Yeaman, Michael C. Whitlock

**Affiliations:** Department of Forest and Conservation Sciences, University of British Columbia, Vancouver, Canada; Biodiversity Research Centre, University of British Columbia, Vancouver, Canada; Department of Biological Sciences, University of Calgary, Calgary, Canada; Department of Zoology, University of British Columbia, Vancouver, Canada

## Abstract

Spatially varying selection promotes variance in allele frequencies, increasing genetic differentiation between the demes of a metapopulation. For that reason, outliers in the genome wide distribution of summary statistics measuring genetic differentiation, such as *F_ST_*, are often interpreted as evidence for alleles which contribute to local adaptation. However, in spatially structured populations, the spread of beneficial mutations with spatially uniform effects can also induce transient genetic differentiation and numerous theoretical studies have suggested that species-wide, or global, adaptation makes a substantial contribution to molecular evolution. In this study, we ask whether such global adaptation affects the genome-wide distribution of *F_ST_* and generates statistical outliers which could be mistaken for local adaptation. Using forward-in-time population genetic simulations assuming parameters for the rate and strength of beneficial mutations similar to those that have been estimated for natural populations, we show the spread of globally beneficial in parapatric populations can readily generate *F_ST_* outliers, which may be misinterpreted as evidence for local adaptation. The spread of beneficial mutations causes selective sweeps at flanking sites, so the effects of global versus local adaptation may be distinguished by examining patterns of nucleotide diversity along with *F_ST_*. Our study suggests that global adaptation should be considered in the interpretation of genome-scan results and the design of future studies aimed at understanding the genetic basis of local adaptation.

## Introduction

Determining the genetic basis of local adaptation is of primary interest to evolutionary biologists, as it provides insights into how natural selection shapes genetic variation. The genetic basis of local adaptation has increasingly been studied using various types of “genome scans,” i.e., methods which are applied across the genome to detect patterns that are expected under the process of local adaptation. Some genome scans look for alleles that are correlated with particular putative selective features of the environment. Others look for genomic regions of particularly high genetic differentiation among populations, based on the fact that local adaptation by definition will increase the frequency of locally beneficial alleles. As a result, local adaptation should cause some increase in the variation among populations in allele frequencies.

One of the most commonly used summary statistics for such genome scans is Wright’s *F_ST_* (or its derivatives), which measures the variance in allele frequencies among the demes of a metapopulation. Since long-term local adaptation promotes variance in allele frequency among demes, regions of the genome containing alleles subject to spatially varying selection should have *F_ST_* values which appear extreme in the genome-wide distribution (Lewontin & Krakauer, 1973). Depending on the strength of selection and the rate of migration, spatially varying selection can result in neutral variants linked to selected alleles also exhibiting signatures expected under local adaptation (Barton & Bengtsson, 1986; Bengtsson, 1985; Petry, 1983). Under this framework, genes or other functional elements linked to *F_ST_* outliers provide researchers with candidate loci to investigate for their role in local adaptation.

However, interpreting the results of genome scans is fraught with difficulties. There are many other reasons aside from local adaptation for why a particular genomic region may have a measure of genetic differentiation that is greater than expected by a particular neutral model. Most genome-scan methods make implicit assumptions about the pattern of population structure and evolutionary history; violation of these assumptions can result in high false positive rates (Lotterhos & Whitlock, 2015).

Additionally, background selection (BGS), the loss of neutral variability at sites linked to those subject to purifying selection (Charlesworth, Morgan, & Charlesworth, 1993), is thought to be ubiquitous in evolution (Comeron, 2017) and in some circumstance can cause elevated *F_ST_* through its effects on genetic diversity (Charlesworth, 1998; Cruickshank & Hahn, 2014; Zeng & Corcoran, 2015; but see Matthey-Doret & Whitlock, 2019). Finally, the spread of adaptive mutations in structured populations affects linked neutral variants and can also influence *F_ST_*.

Adaptation via uniformly selected alleles—referred to here as “global adaptation”—can also affect patterns of *F_ST_*. The spread of uniformly advantageous alleles causes a reduction in nucleotide diversity, or heterozygosity, at linked neutral sites (Maynard Smith & Haigh, 1974), a process referred to as a selective sweep. During the course of a selective sweep in structured populations, genetic differentiation, or *F_ST_*, at linked sites may become elevated (Barton, 2000; Bierne, 2010; Feder et al., 2019; Kim & Maruki, 2011; Santiago & Caballero, 2005; Slatkin & Wiehe, 1998). In parapatric populations for example, advantageous mutations are likely to be temporarily at higher frequencies in the region where they originated, before they migrate, establish, and fix in other populations. This introduces a lag, during which sites linked to the selected locus exhibit increased genetic differentiation. As the beneficial allele establishes and spreads in the other deme, the differentiation decays (Kim & Maruki, 2011; Slatkin & Wiehe, 1998). In addition, recombination events which occur during the course of the sweep can introduce the beneficial allele onto different genetic backgrounds in different demes, which subsequently hitchhike to high frequencies, generating differentiation. Once the beneficial allele fixes in such cases, twin peaks of differentiation may be left around the selected site which is then eroded by subsequent migration (Bierne, 2010). Thus, if global selective sweeps in structured populations are common, they may influence the genomic patterns of *F_ST_*.

How common are such global selective sweeps? In eukaryotes, the best information on the process of global adaptation is from *Drosophila melanogaster*. An estimate of the rate of selective sweeps in *D. melanogaster* can be obtained as follows: *D. melanogaster* and *D. simulans* began to diverge an estimated 14 million generations ago (Obbard et al., 2012). In that time, *D. melanogaster* has accumulated 0.0067 substitutions/bp at nonsynonymous sites (Begun et al., 2007). The proportion of nonsynonymous substitutions driven by positive selection (*α*) in *D. melanogaster* has been estimated in numerous studies to be ~0.5 (Begun et al., 2007; Elyashiv et al., 2016; Eyre-Walker & Keightley, 2009; Messer & Petrov, 2013). There are approximately 15 Mbp of nonsynonymous sites in the *D. melanogaster* genome, calculated as 2/3 of the sites in protein-coding regions (based on *D. melanogaster* genome build 6.28; downloaded from FlyBase FB2019_03; Thurmond et al., 2019). Taken together, this suggests that an estimated (0.5 x 15 x 0.0067)/14 = 0.0035 nonsynonymous substitutions have occurred each generation since *D. melanogaster* split with *D. simulans*. Put another way, substitutions of advantageous alleles have occurred approximately every 280 generations in *D. melanogaster*. Thus, if advantageous mutations take longer than 280 generations to fix, multiple sweeps will be going on at any point in time. With knowledge of the effective population size (*N_e_*) and selection coefficients for beneficial mutations (*s_a_*), expected fixation times can be calculated (Ewens, 1979). It has been estimated that advantageous nonsynonymous mutations in *D. melanogaster* have scaled effects on fitness of *2Nesa* = 250 (Campos et al., 2017). Assuming a historical *N_e_* for *D. melanogaster* of 10^6^, advantageous mutations at nonsynonymous sites in *D. melanogaster* would take ~97,500 generations to fix and spend ~22,200 generations at intermediate frequencies (i.e. between 0.2 and 0.8), assuming a panmictic population. Taken together, these calculations suggest that at any point in time, *D. melanogaster* has approximately 22,200/280 = 80 incomplete sweeps of alleles that will ultimately reach near fixation. This back-of-the-envelope calculation is obviously quite rough; the values used are estimates which may be biased and there is likely a distribution of fitness effects for new beneficial mutations. If anything, this likely underestimates the number of ongoing sweeps, as population structure prolongs the time to fixation above the expectation under panmixia (Whitlock, 2003) and beneficial mutations may occur at sites outside protein-coding regions (for example, in the untranslated region of genes). Nevertheless, this approximation gives an idea of the order of magnitude of the number of ongoing sweeps.

If there are numerous ongoing sweeps in a population, as suggested by the *Drosophila* calculation, some incomplete global sweeps may induce genetic differentiation and be detected when scanning the genome for local adaptation. In this study, we analyse the genomic landscape of genetic differentiation in structured populations subject to recurrent selective sweeps. In particular, we focus on the rate at which selective sweeps of globally beneficial mutations can generate ephemeral *F_ST_* peaks and thus resemble local adaptation in genome scans. We simulate structured populations undergoing recurrent adaptation and perform genome scans on the resulting data. Our results suggest that advantageous mutation parameters compatible with published rates of adaptive evolution can readily generate *F_ST_* outliers in genome scans. We also find that nucleotide diversity in regions containing *F_ST_* outliers help distinguish local versus global adaptation. Our study highlights that global adaptation may be a common cause of *F_ST_* outliers and should be considered when designing studies and interpreting the results of genome scans.

## Methods

### Two-deme simulations

We modelled global adaptation in parapatric populations using forward-in-time simulations in *SLiM* 3.2 (Haller et al., 2019; Haller & Messer, 2019). We simulated an initial Wright-Fisher population of *N* = *N_e_* = 10,000 diploid individuals (unless otherwise stated) which is later split into two equally sized demes of 5,000 individuals each. Symmetrical migration occurred between the demes with probability m. Simulated genomes consisted of 20 “gene-like regions” of 5,000 bp each separated by 100,000 bp of neutral sequence. Advantageous mutations occurred in the “gene-like regions” at a rate of *μp_a_* per generation and their spatially uniform fitness effects were drawn from an exponential distribution with mean 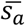. We roughly based our simulations on natural *Drosophila melanogaster* populations, where the effective mutation and recombination rates, *4Nμ* and *4Nr* respectively, are typically estimated to be around 0.01 (Chan et al., 2012; Langley et al., 2012). Using the tree-sequence option in *SLiM*, we recorded the coalescent histories of our simulations every 40,000 generations after the initial population split for 400,000 generations, and thus each simulation replicate gave us a total of 10 quasi-independent datasets. Neutral mutations were overlaid upon simulated genealogies at a rate of *μ* = 2.5 x 10^-7^ per bp per generation. Recombination occurred at a uniform rate of *r* = 2.5 x 10^-7^ per bp per generation. For a given set of selection and migration parameters (Table S1), we performed 200 replicate simulations. In lieu of performing extensive burn-in, we used the ‘recaptitation’ feature of *PySlim*, which simulates the coalescent history of the population before the start of simulations using *msprime* (Kelleher et al., 2016). In order to minimise bias that may arise from grafting forward-in-time genealogies to those simulated under the coalescent, the forward component of our simulations included 1,000 generations of neutral evolution before the initial population split.

In order to model local adaptation, we performed additional simulations which were identical to those described above except for the following: We introduced a single copy of an antagonistically pleiotropic allele in the centre of a 600 kbp chromosome 9,000 generations after the initial population split into two demes. The selected allele conferred a heterozygotic fitness of 1 + *s_a_* in its deme of origin, but 1 / (1 + *s_a_*) in the other. Simulations where the selected allele was lost were discarded.

### Analysis of simulated data

From our simulation data, samples of 25 diploid individuals were drawn from each deme and used to generate VCF files. Weir & Cockerham’s (1984) estimator of *F_ST_* was then calculated for individual SNPs or from analysis windows of 10,000 basepairs using VCFtools (Danecek et al., 2011). For each analysis window we also calculated nucleotide diversity in the metapopulation as a whole (*π_T_*).

In order to classify analysis windows or SNPs as outliers, we used the distribution of *F_ST_* from neutral simulations. In all aspects, the neutral simulations were identical to those modelling global adaptation, except that the *p_a_* parameter was set to 0. For each migration rate we tested (Table S1), we performed a set of 200 neutral simulation replicates and from them obtained the distribution of *F_ST_*, either per analysis window or per SNP. Any analysis window or SNP with an *F_ST_* value beyond the 99.999th percentile of the distribution was considered an outlier. For the purpose of comparing parameter sets (Table S1), we only considered the analysis windows centred on the simulated “gene-like regions”. For each parameter set, we examined *F_ST_* across 40,000 “gene-like regions”, we used the 99.999th percentile of the distribution from neutral loci as a conservative *F_ST_* cut-off. With this cut-off, we expect 0.4 outliers per parameter set in the purely neutral case.

We further analysed the per SNP *F_ST_* values using a version of the top-candidate test of (Yeaman et al., 2016). We calculate the average proportion of outlier SNPs present in 10,000 bp regions from the neutral simulations, as 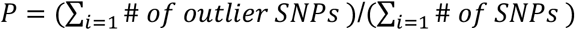, summing over *i* regions. The number of outlier SNPs present in 10,000 bp analysis windows centred on the gene-like regions was then treated as a binomial variable with probability *P*. The “gene-like” regions with a number of outlier SNPs greater than the 99.99th percentage point of the binomial distribution are classified as top-candidates. We note that *P* should be considered an index, rather than an accurate probability, due to the non-independence among SNPs within a 10,000 bp region, which violates the assumptions of the binomial distribution.

### Data availability

All analysis scripts and simulation configuration files are available at https://github.com/TBooker/GlobalAdaptation.

## Results

### Ongoing and recent selective sweeps of globally beneficial mutations confound genome scans

In spatially-structured populations, selective sweeps of globally beneficial alleles may influence the landscape of genetic differentiation. The Manhattan plot in Figure 1 shows a typical sliding-window genome scan for *F_ST_* performed on data from a simulated parapatric population. For the purposes of demonstration the simulations shown in Figures 1 and 2 were performed using *N* = 2,000 diploid individuals (1,000 per deme). The simulated genome consisted of 500 Mbp of simulated sequence, with 25 Mbp of functional sites. The number of functional sites was chosen to approximately reflect the total number of nonsynonymous sites and sites in the untranslated regions of protein-coding genes in *D. melanogaster*. Advantageous mutations occurred at functional sites with fitness effects and rates similar to those that have been estimated for *D. melanogaster* (Campos et al., 2017). This particular genome scan identified two regions containing *F_ST_* outliers, defined as those with *F_ST_* greater than the 99.999th percentile of values from neutral simulations (given our cut-off value, we expected an average of 0.05 outlier regions under neutrality). When sampling the same population at different time points, a similar number of *F_ST_* outliers are observed but they are in different genomic locations (Figure S1). Note that at some time points no outliers will be observed (Figure S1). If one performed a genome scan on real data and had observed outliers such as those in Figure 1, it would be tempting to interpret them in terms of spatially varying selection and local adaptation, when in fact they were caused by the transient differentiation induced by the fixation of globally beneficial alleles.

**Figure 1.**
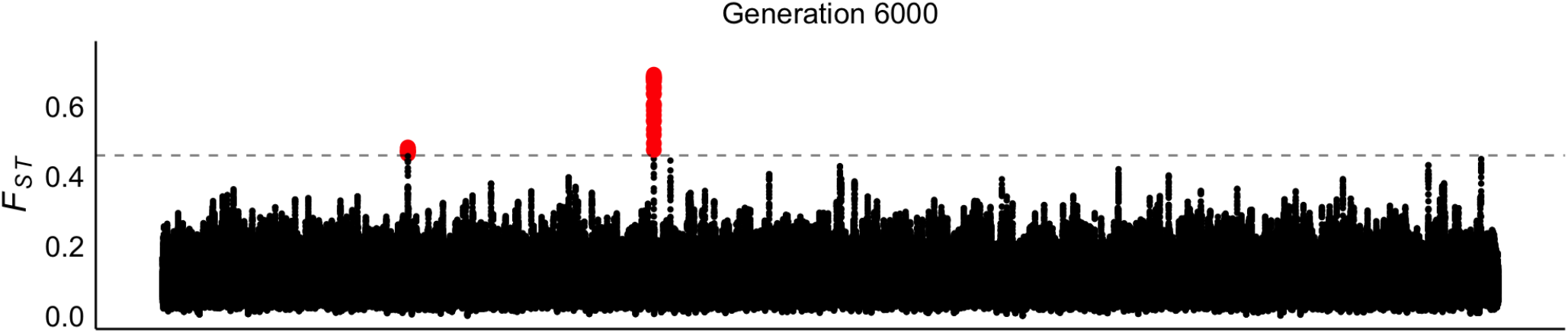
Manhattan plot of *F_ST_* calculated between parapatric populations subject to global adaptation. Sliding windows of 10,000 bp with a step size of 500 bp were used. The horizontal grey line indicates the 99.999th percentile of *F_ST_* from neutral simulations. Simulation parameters were *N* = 2,000 diploid individuals, 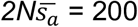, *p_a_* = 0.0001 and *Nm* = 1.

The fixation of globally beneficial mutations may generate *F_ST_* outliers by at least two processes. Firstly, beneficial mutations may spread to high frequency in their demes of origin with a lag before migrating and establishing in other demes. If this period of lag is substantial, it may correspond to an ephemeral increase in *F_ST_*. Secondly, recombination may move beneficial alleles onto different genetic backgrounds in different demes which may then be driven to high frequency during the sweep. Thus, in the wake of the sweep, there may be elevated *F_ST_* in regions surrounding selected loci, which is then eroded by subsequent migration (Bierne, 2010). *F_ST_* peaks generated by both processes are shown in Figure 2. For example, the pink lines in Figure 2 show the change in allele frequency over time for one particular beneficial mutation and the associated *F_ST_*. The beneficial mutation occurs in one deme and sweeps to high frequency before it establishes and spreads in the other (Figure 2A). During this period of lag, *F_ST_* in the region around the selected site is elevated, but drops off once the mutation reaches high frequency in the second deme (Figure 2B). The yellow lines in Figure 2 show that elevated *F_ST_* can persist once a beneficial mutation has gone to fixation, and this may occur as a result of the process described by (Bierne, 2010). For the purpose of visualisation, we display the allele frequency and associated *F_ST_* profiles for seven beneficial alleles in Figure 2, but many more may be segregating in the population as a whole at any one time generating a heterogeneous landscape of differentiation as in Figure 1.

**Figure 2.**
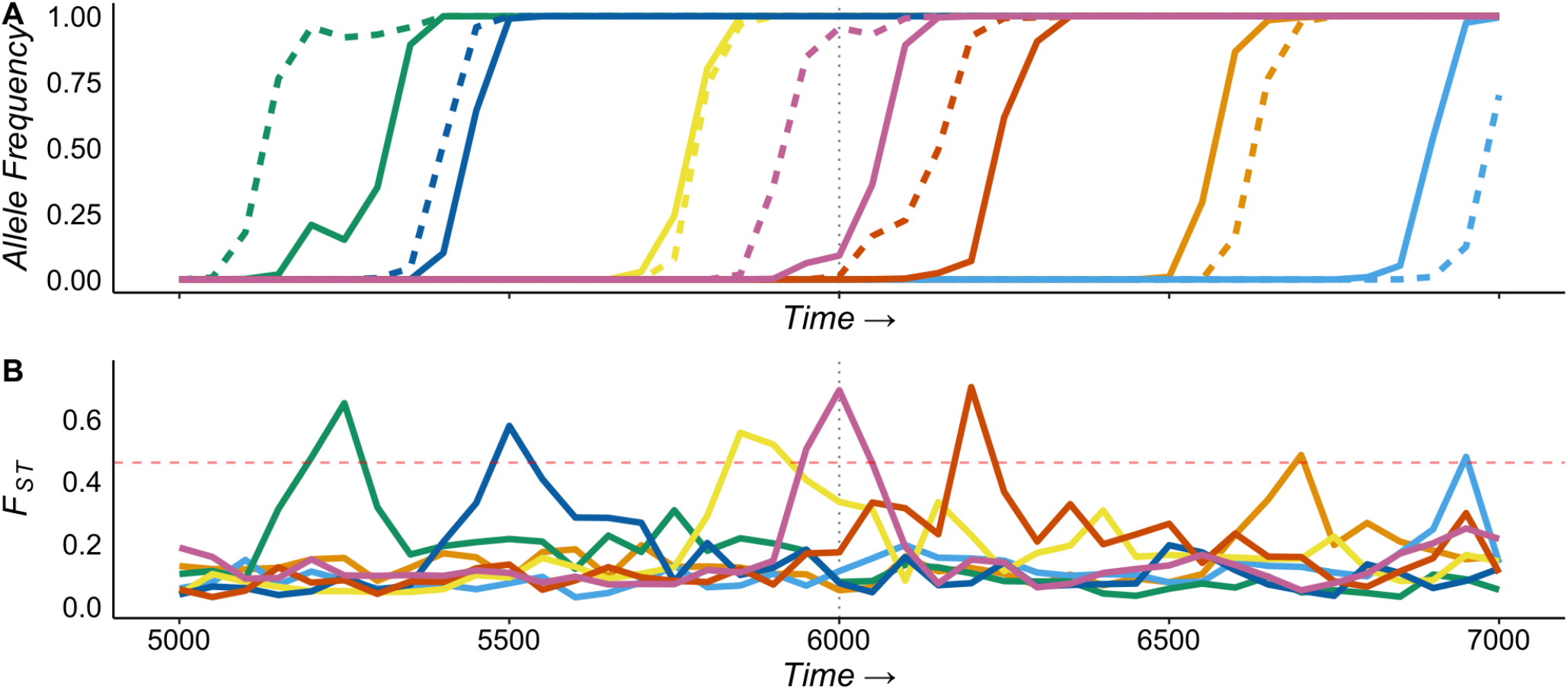
Selective sweeps of globally beneficial alleles can generate ephemeral *F_ST_* peaks. (A) the allele frequency of beneficial mutations in deme 1 (solid lines) and deme 2 (dashed lines). (B) *F_ST_* over time for 10,000 bp analysis windows containing beneficial alleles. The vertical line indicates the time at which the genome scan shown in Figure 1 was performed. The dashed horizontal line indicates the 99.999th percentile of *F_ST_* from neutral simulations. Simulation parameters, *N* = 2,000 diploid individuals, 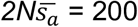, *p_a_* = 0.0001, *Nm* = 1.

### The frequency of *F_ST_* outliers generated by global adaptation

Global adaptation can influence the genomic landscape of differentiation (Figures 1 and 2), but the extent to which it does depends on the rate and selective effects of beneficial mutations. With a greater number of mutations sweeping to fixation at any one time, there will be a greater chance of observing *F_ST_* peaks driven by global adaptation. We simulated parapatric populations subject to advantageous mutations with a range of DFEs which corresponded to α values from 0.02 to 0.61 (Table S1). The range of α values exhibited by our simulated populations is consistent with the range estimated by Galtier (2016) for numerous eukaryotic species. We measured the frequency of *F_ST_* outliers driven by globally beneficial mutations by calculating the proportion of 10,000 bp analysis windows with *F_ST_* greater than the 99.999th percentile of values from neutral simulations. Figure 3 shows that the proportion of analysis windows containing outliers increases with the rate and strength of advantageous mutations in parapatric populations. Comparing the same DFE, but under different migration rates, Figure 3 also shows that outliers driven by global adaptation are more frequent when *Nm* = 1 than when *Nm* = 10. This is expected, as the delay in the fixation time for beneficial alleles in structured populations decreases with both *m* and *s_a_* (Kim & Maruki, 2011) and *F_ST_* peaks may occur as a result of this period of lag. However, not all outliers that we detected were linked to ongoing sweeps (Figure S2), suggesting that the process described by Bierne (2010) may substantially contribute to heterogeneity in *F_ST_* across our simulated genomes. Qualitatively similar results were found using a SNP-based genome-scan method (Figure S3). However, the proportion of outlier regions detected using the SNP-based method was lower than when using analysis windows (Figure S3), suggesting that the different approaches have different sensitivities to the effects of global adaptation.

**Figure 3.**
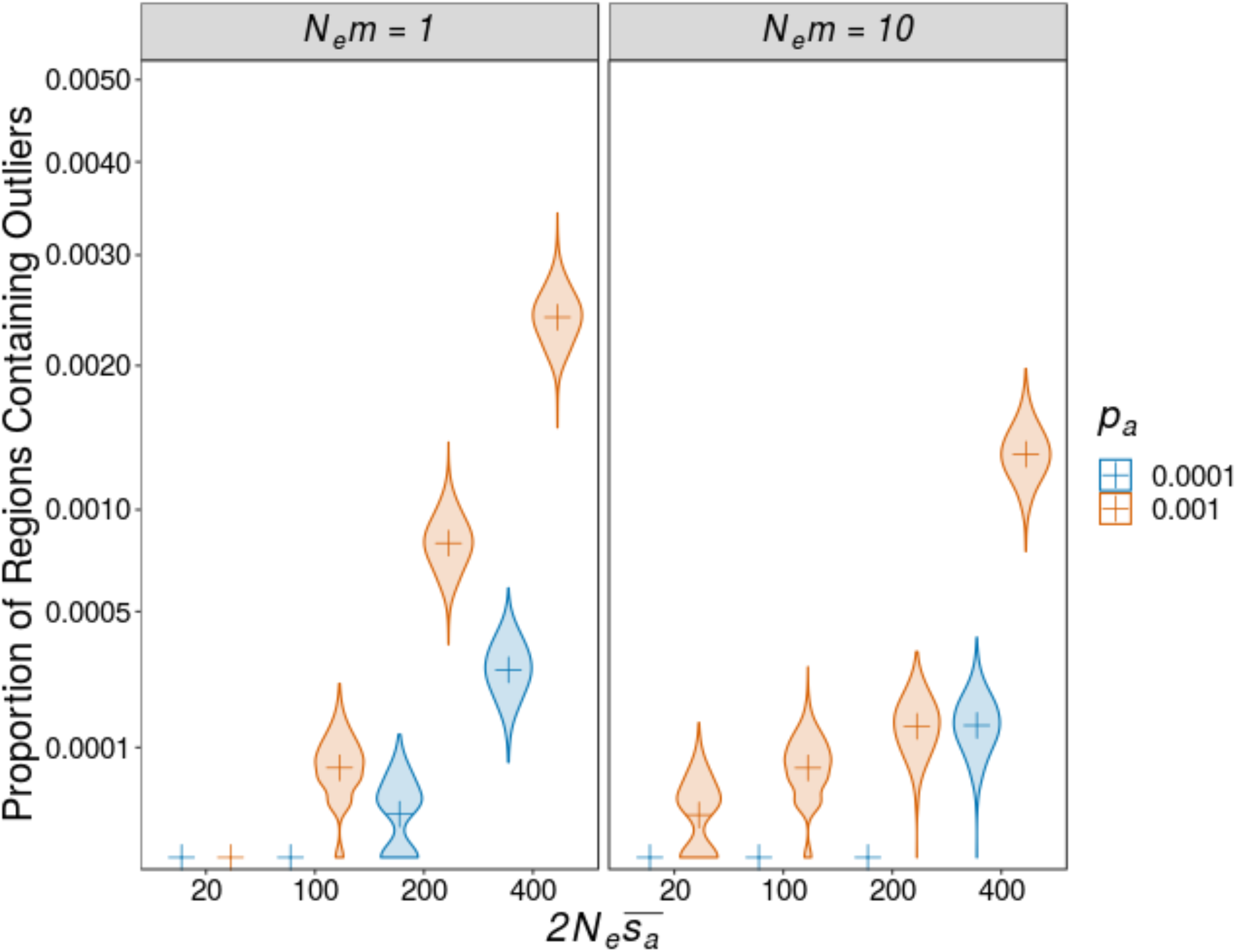
The proportion of analysis windows that contain *F_ST_* outliers in parapatric populations. Allele-frequency weighted *F_ST_* was calculated for 10,000 bp analysis windows centred on simulated “gene-like regions”. Plusses indicate the point estimate and violins indicate the distribution of 1,000 bootstraps samples from 2,000 simulated datasets.

In continuously distributed populations, globally advantageous mutations spread in a wave-like fashion (Fisher, 1937; Kolmogorov et al., 1937) which can generate allele frequency clines at linked sites (Barton, 2000; Barton et al., 2013). When comparing populations from different points in a continuously distributed species’ range, allele frequency clines generated by the spread of globally beneficial mutations could resemble local adaptation, in a manner similar to the two-deme case studied above. We approximated adaptation in continuously distributed populations by simulating a one-dimensional stepping-stone model with many demes. (See Supplemental materials for a description of the methods for these simulations.) In these simulated populations, the number of *F_ST_* outliers at neutral sites linked to selected loci were positively related to the rate and average strength of advantageous mutations (Figure S4). When contrasting points in a continuous range, the probability of an ongoing sweep separating the two increases with physical distance (Appendix 1), and accordingly, so does the number of *F_ST_*, outliers (Figure S4). As advantageous mutations spread through the population, recombination decouples them from linked neutral sites, decreasing the frequency of *F_ST_* outliers (Figure S4).

### Distinguishing local from global adaptation

Long-term local adaptation results in a buildup of linkage disequilibrium and neutral genetic diversity in genomic regions surrounding alleles subject to spatially varying selection (Charlesworth et al., 1997; Petry, 1983). In structured populations, selective sweeps of globally beneficial alleles reduce genetic diversity in surrounding regions across the metapopulation as a whole (Barton, 2000). The different effects that global versus local adaptation have on linked diversity may, therefore, provide a means to distinguish them. Figure 4 shows *F_ST_* against the ratio *π_T_/π_0_*, where *π_T_* is nucleotide diversity across the metapopulation and *π_0_* is the expectation for a panmictic population under neutrality. In the absence of selection, genetic drift may generate strikingly large *F_ST_*, but in such cases, the underlying nucleotide diversity is not expected to systematically vary from the genomic background (Figure 4). As expected, *F_ST_* outliers generated under both global and long-term local adaptation have opposite effects on total nucleotide diversity (Figure 4). This suggests that examining the patterns of diversity surrounding *F_ST_* outliers provides a way to infer which kind of selective processes have occurred. Indeed, Bierne, (2010) showed that the patterns differentiation and heterozygosity at loci surrounding an *F_ST_* outlier detected in the marine mussel *Mytilus edulis* were better predicted by a model of global adaptation than a model of local selection.

**Figure 4.**
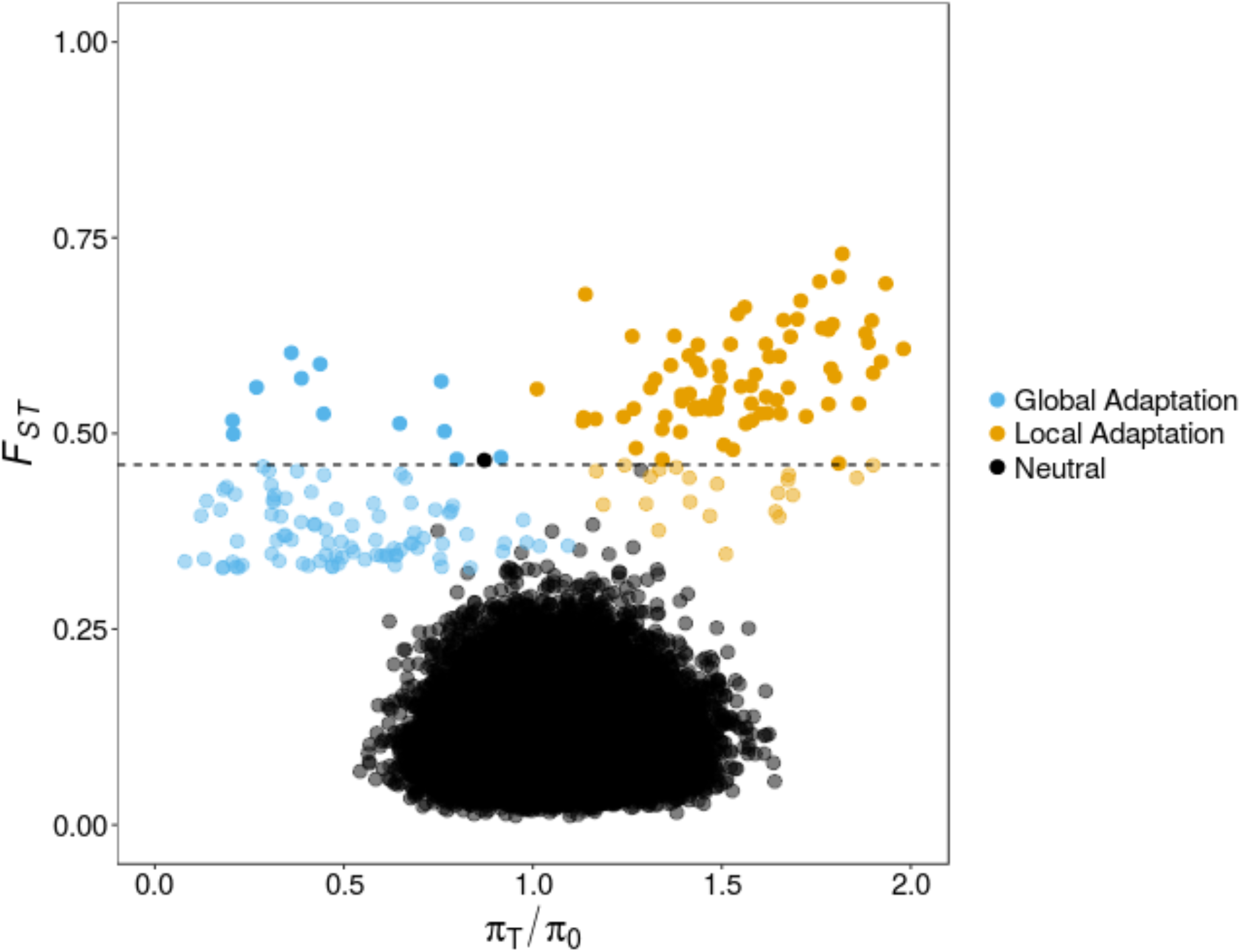
Scatterplot of *F_ST_* against relative nucleotide diversity (*π_T_/π_0_*) for outliers driven by global versus local adaptation compared with neutrality. In all cases, results for analysis windows of 10 Kbp are shown. In the case of local adaptation, an antagonistically pleiotropic allele with *2Ns_a_* = 200 was simulated. The points labelled ‘global adaptation’ are from the analysis windows, centred on genes, with the top 100 values from simulations with 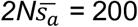 and *p_a_* = 0.0001 shown. In all simulations *Nm* = 1.

### The number of incomplete range-wide selective sweeps

We have shown that global adaptation in structured populations may be detected as *F_ST_* outliers in genome scans. In both the cases of parapatric and stepping-stone populations, outliers can be driven by incomplete selective sweeps (Figures S2 and S4), so understanding the expected number of incomplete sweeps for a given distribution of fitness effects would be useful. Fixation times for beneficial alleles are longer in structured populations than under panmixia by an amount that is proportional to 1 - *F_ST_* (Whitlock, 2003). Thus, it seems reasonable to conclude that, all else being equal, the number of advantageous mutations spreading to fixation at any one time will be greater for a structured population than under panmixia. We therefore modelled the number of incomplete selective sweeps under panmixia to obtain a lower bound for the expected number in structured populations (Appendix 2).

Under a variety of DFEs, a large number of incomplete sweeps are expected at any one time in panmictic populations (Figure 5A). For the same rate of beneficial mutations, the number of incomplete sweeps increases with the mean of the DFE (Figure 5A). As the mean of the DFE increases beyond 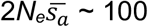, however, the fixation probability and time beneficial alleles spend at intermediate frequencies balance out and the number of incomplete sweeps does not increase (Figure 5A). As expected, the number of incomplete sweeps in our simulated parapatric populations exceeded the expectation under panmixia in all cases (Figure 5A). Estimates of the DFE for advantageous mutations have only been obtained for a handful of organisms, but the proportion of adaptive substitutions (*α*) has been estimated in numerous species to be around 0.5 (e.g. Galtier, 2016). Figure 5A shows that numerous DFEs for beneficial mutations result in many incomplete sweeps, and Figure 5B shows that the same DFEs give rise to *α* values consistent with those that have been published. Taken together, these results suggest that the number of incomplete sweeps in natural populations may be quite large, and if that is the case, the chances of observing *F_ST_* outliers when performing studies of local adaptation is as well.

**Figure 5.**
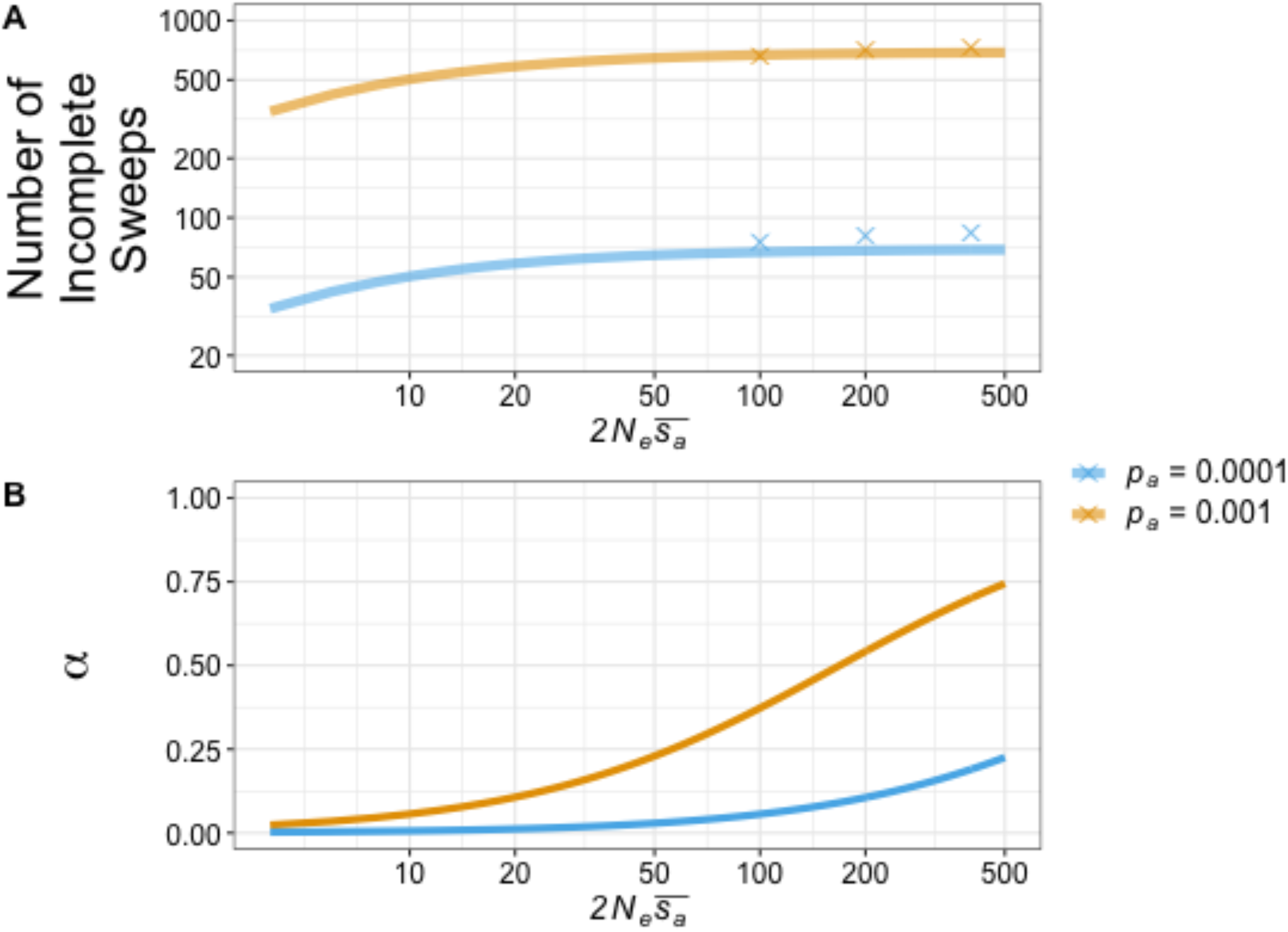
The number of incomplete selective sweeps and the proportion of adaptive substitutions (α) under exponential DFEs for advantageous mutations. A) The solid lines show the expected number of incomplete selective sweeps in panmictic populations, calculated from Equation A5, the crosses are from parapatric populations simulated under *Nm* = 1. B) The proportion of substitutions driven by positive selection. The solid lines were obtained using Equation A6. Parameters not specified in the plot, *N_e_* = 10,000, *4N_e_μ* = 0.01, *η_a_* = 2.5 x 10^7^.

## Discussion

In the search for loci underpinning local adaptation, *F_ST_* outliers driven by global adaptation are a source of false positives. However, such outliers are not statistical artefacts; rather, they are highly differentiated regions of the genome, so methods that are aimed at detecting local adaptation by examining outliers may be sensitive to the effects of global adaptation. In this study, we obtained the neutral distribution of *F_ST_* from simulations with identical demography to those modelling selection, allowing us to define outliers using stringent cut-off values. In most real-world cases, however, population structure and demography are not fully understood, so determining outliers in this way is not possible. Methods for analyzing results from genome scans may examine the tails of the empirical distribution of *F_ST_* (e.g. Reinhardt et al., 2014) or look for genomic regions with an excess of highly differentiated SNPs (e.g. Mateo et al., 2018; Yeaman et al., 2016). In either of these cases, global adaptation could be mistaken for local adaptation, because it can cause an ephemeral increase in genetic differentiation (Figure 2). There are genome scan methods for detecting local adaptation that are more sophisticated than the comparatively simple *F_ST_* scans we implemented in this study, for example *FDist2, BayScan* and *OutFlank* (Beaumont & Nichols, 1996; Foll & Gaggiotti, 2008; Whitlock & Lotterhos, 2015). However, even such methods may be misled by global adaptation because the differentiation it induces can be highly similar to that expected under long-term local adaptation (Figure 4).

### Study design will influence how global adaptation affects genome scans

The confounding effects of global adaptation discussed in this study arise because populations may be sampled along a linear array that is parallel to a possible path taken by a globally sweeping mutation. This would be analogous to sampling along a coastline or a river valley, where a single axis determines both the spatial patterning of gene flow and environmental change. While we did not explicitly study genotype-environment association analyses (e.g. Bayenv; Coop et al., 2010), global adaptation would also affect such studies if the environmental gradient in question has a simple spatial structure that covaries linearly with demography. The simplest solution to this problem would be to sample populations across multiple replicates of a given transition in environment, each distributed in different parts of a species range, such that it would be exceedingly unlikely that a given globally sweeping mutation would simultaneously be constricted in its spatial range to populations of a given type of environment (as per Figure 1, Lotterhos & Whitlock, 2015). Unfortunately, this will not be possible in many species, if the environment of interest varies only along a single spatial dimension.

As discussed above, it may sometimes be possible to distinguish local adaptation from global sweeps based on the patterns of within-population nucleotide diversity at flanking loci. Whereas many kinds of selection can cause a reduction in nucleotide diversity at flanking loci, only prolonged migration-selection balance would cause both a peak in *F_ST_* and an increase in within-population nucleotide diversity at flanking loci (Petry, 1983). Unfortunately, the lack of such an increase in nucleotide diversity is not evidence against local adaptation, as it would be difficult to distinguish young locally adapted alleles and the effects of global adaptation.

### Models of adaptation and population structure

In this study, we have assumed that positive selection acts on *de novo* mutations, causing “hard” selective sweeps, but other modes of adaptation may be frequent in nature. When positive selection acts on standing variation, “soft” selective sweeps can occur (Hermisson & Pennings, 2017) and these may be common (Garud et al., 2015; Schrider & Kern, 2017). For instance, Reinhardt et al. (2014) performed genome scans on natural *D. melanogaster* populations distributed along the Eastern seaboards of Australia and North America. They found *F_ST_* outliers common to both populations, suggesting that selection may have acted on shared standing genetic variation. In spatially extended populations, multiple copies of the same allele may arise through independent mutational events causing parallel selective sweeps (Paulose, Hermisson, & Hallatschek, 2019; Ralph & Coop, 2010; Ralph & Coop, 2015). In the cases of both soft and parallel sweeps in structured populations, the extent to which global adaptation will influence *F_ST_* likely depends upon the differentiation in the region of the selected locus at the onset of selection. If soft sweeps have contributed substantially to estimates of a, the back-of-the-envelope calculation we developed here for the number of simultaneous sweeps may somewhat overestimate the number of loci with substantial *F_ST_* peaks.

Ultimately, whether global adaptation induces *F_ST_* outliers in subdivided populations will depend on the rates of adaptation and gene flow. In a study of African populations of *D. melanogaster*, Vy et al. (2017) found evidence that 37 loci across the entire genome were currently experiencing selective sweeps. However, the methods they used were powered to detect very strong selection (*2N_e_s_a_* > 2,000), so it seems likely that alleles with more modest effects on fitness may have been missed by their approach. In our introduction, using a coarse calculation, we estimated the number of incomplete sweeps at nonsynonymous sites in *D. melanogaster* to be around 80. If the number of ongoing sweeps in natural populations is of similar magnitude, then there may be ample opportunity for global adaptation to induce *F_ST_* outliers and confound genome scans. Finally, environments vary over space, so geographically distant populations may be subject to distinct biotic and abiotic environments. Studies may sample geographically distant populations in order to maximise the extent by which local adaptation shapes patterns of genetic differentiation. However, increasing the distance between sampled demes, which for species with limited dispersal is analogous to reducing the migration rate in an island model, increases the chance that global adaptation induces *F_ST_* outliers (Figure S4).

## Conclusions

In this study, we have shown that recurrent global adaptation in structured populations can give rise to *F_ST_* peaks, which may frequently be detected as outliers in genome scans (Figure 1). The idea that global adaptation influences the genomic landscape of differentiation is not novel to this study; indeed it has been invoked to explain patterns of differentiation in natural populations before (Bierne, 2010). However, we have shown that, under rates of adaptation consistent with results from population genomic studies in numerous animal and plant species (e.g. Galtier, 2016; Hodgins et al., 2016; Williamson et al., 2014), *F_ST_* peaks driven by globally beneficial mutations are likely to be a common feature of the landscape of differentiation in structured populations. For that reason, global adaptation should be considered in the design of future studies and in the interpretation of genome scan results.

## Acknowledgments

We wish to thank Andréa Thomaz and members of the Whitlock lab for helpful discussions. This paper is part of the CoAdapTree project which is funded by Genome Canada (241REF), Genome BC and 16 other sponsors (http://coadaptree.forestry.ubc.ca/sponsors/). MCW is supported by a Discovery Award from NSERC. SY is supported by a Discovery Award from NSERC and a research chair from Alberta Innovates.

## Author Contributions

T.R.B, S.Y. and M.C.W. designed the study. T.R.B. performed all simulations and analyses. T.R.B. wrote the article with input from S.Y. and M.C.W.

## Supplementary Material

### Supplementary Methods

#### Two-locus Stepping-Stone Simulations

We approximated the process of global adaptation in continuous space by simulating populations structured according to a one-dimensional stepping stone model. Stepping stone simulations consisted of a linear array of *k* = 500 demes arranged next to one another, with *N* = 5,000 haploid individuals per deme. Nearest-neighbour dispersal between demes occurred with probability *m* = 0.666 (0.333 in either direction). The genomes of each haploid individual consisted of two biallelic loci *A/a* and *B/b*. Individuals possessing the *a* allele had a fitness of 1 and those bearing the *A* allele had a selective advantage of 1 + s in all demes. The *B/b* locus was strictly neutral. Our simulations tracked the frequencies of the four possible genotypes (*AB, Ab, aB* and *ab*) through time. Recombination between the *A/a* and *B/b* loci occurred with probability *c*. Each generation, genotype frequencies were affected by the deterministic forces of selection, recombination and migration (in that order) and multinomial sampling was then used to create offspring, simulating genetic drift.

In order to achieve an equilibrium distribution of neutral allele frequencies across our simulated metapopulations, a period of burn-in was performed. However, our stepping-stone simulations consisted of 2.5 x 10^6^ individuals and in such populations, simulating the mutation and subsequent equilibration of neutrally evolving alleles would take an unfeasibly long time. In order to reduce simulation time, we provided a starting point for the neutral burn-in by assuming a correlation in allele frequencies between adjacent demes of *r* = 0.99. The value of *r* we used was roughly based on results for high migration rates in the stepping-stone model given in Kimura & Weiss (1964). For a given simulation, we generated allele frequency profiles as follows. A deme was selected at random and the frequency of the *b* allele (*p_b_*) was set using a draw from a uniform distribution *U*(0, 1). Moving in both directions away from the initial deme, correlated allele frequencies were sampled using *r p_b_*(1 + *N*(0, 1 - *r*^2^)). Correlated allele frequencies were chosen until all demes have been assigned an initial value. Following the initial assignment of allele frequencies, we ran individual replicates for 100,000 generations of drift, discarding runs if the variant at the neutral locus was lost. We performed a total of 100,000 neutral simulations, recording the final allele frequency in each deme.

To simulate the spread of advantageous mutations with varying selection coefficients, we sampled a neutral simulation from the 100,000 replicates we performed and introduced a single copy of an advantageous mutation into a randomly selected deme. For all values of *s_a_* from 0.0001 to 0.1 (in increments of 0.0001), we performed 200 replicate simulations. We discarded runs where the advantageous allele was lost.

Under a given model of the DFE, we simulated the process of advantageous mutations occurring in our stepping-stone simulations as follows. Each generation, we draw a number of new advantageous mutations that occurred in the population as a whole, Poisson-distributed with mean *U_a_*, and assign a selection coefficient to each according to draws from the DFE. Each mutation fixes with probability 2*s_a_*, and for those that do we sample a two-locus simulation with the appropriate selection coefficient. This process is repeated for 100,000 generations. At generation 99,999 we calculate *F_ST_* for neutral loci between pairs of demes using the formula for haploids from (Weir, 1994). This procedure was repeated 30 times for each combination of *U_a_* and 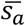.

**Table S1.** Parameters of the DFE for advantageous mutations used in this study and the corresponding *α* values under a model of parapatry. For the purposes of determining *α*, a fraction 1 - *p_a_* of sites are subject to a gamma DFE for harmful mutations with shape parameter 0.3 and scale/mean 2*Ns_d_* = 200, where *s_d_* is the selection coefficient against deleterious alleles. The DFE for harmful mutations was estimated for *Drosophila melanogaster* by Loewe & Charlesworth (2006).

| $2Ns_a$ | $p_a$ | $\alpha$ | |
| --- | --- | --- | --- |
| | | $N_e m = 1$ | $N_e m = 10$ |
| 400 | 0.0001 | 0.237 | 0.238 |
|  | 0.001 | 0.605 | 0.609 |
| 200 | 0.0001 | 0.165 | 0.155 |
|  | 0.001 | 0.503 | 0.503 |
| 100 | 0.0001 | 0.096 | 0.093 |
|  | 0.001 | 0.393 | 0.392 |
| 20 | 0.0001 | 0.020 | 0.020 |
|  | 0.001 | 0.150 | 0.151 |

$2Ns_a$ - The mean of the exponential distribution of fitness effects for advantageous mutations
$p_a$ - The proportion of new nonsynonymous mutations which are advantageous
$\alpha$ - The proportion of substitutions at functional sites driven by positive selection

**Figure S1.**
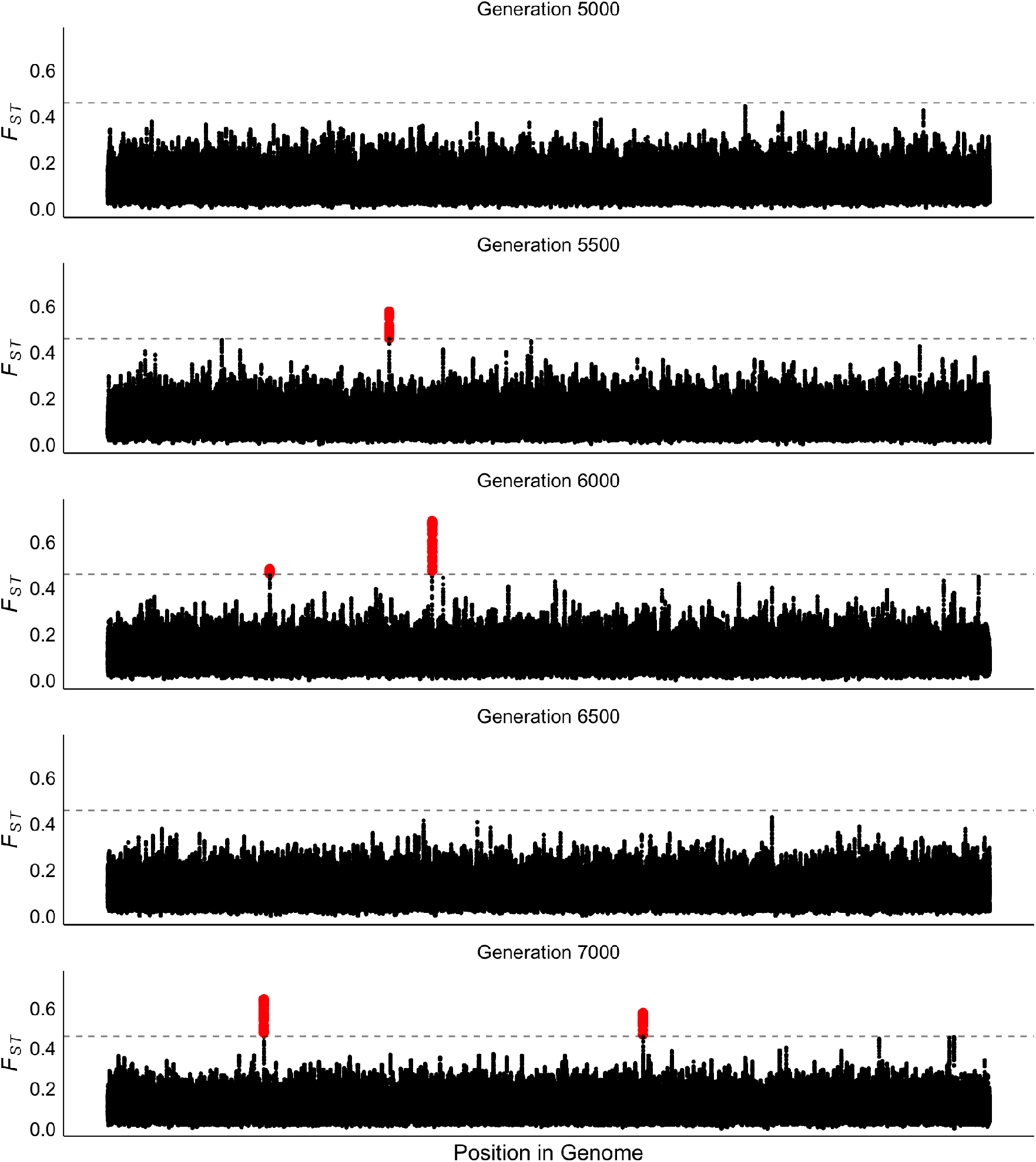
Manhattan plots of *F_ST_* calculated between parapatric populations subject to global adaptation at several time points. *F_ST_* was calculated in sliding windows of 10,000 bp with a step size of 500 bp. The dashed horizontal line shows the 99.999th percentile of *F_ST_* at neutral sites. Simulation parameters, *N* = 2,000 diploid individuals, 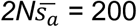, *p_a_* = 0.0001, *Nm* = 1. The central panel, Generation = 6000, is Figure 1 in the main text.

**Figure S2.**
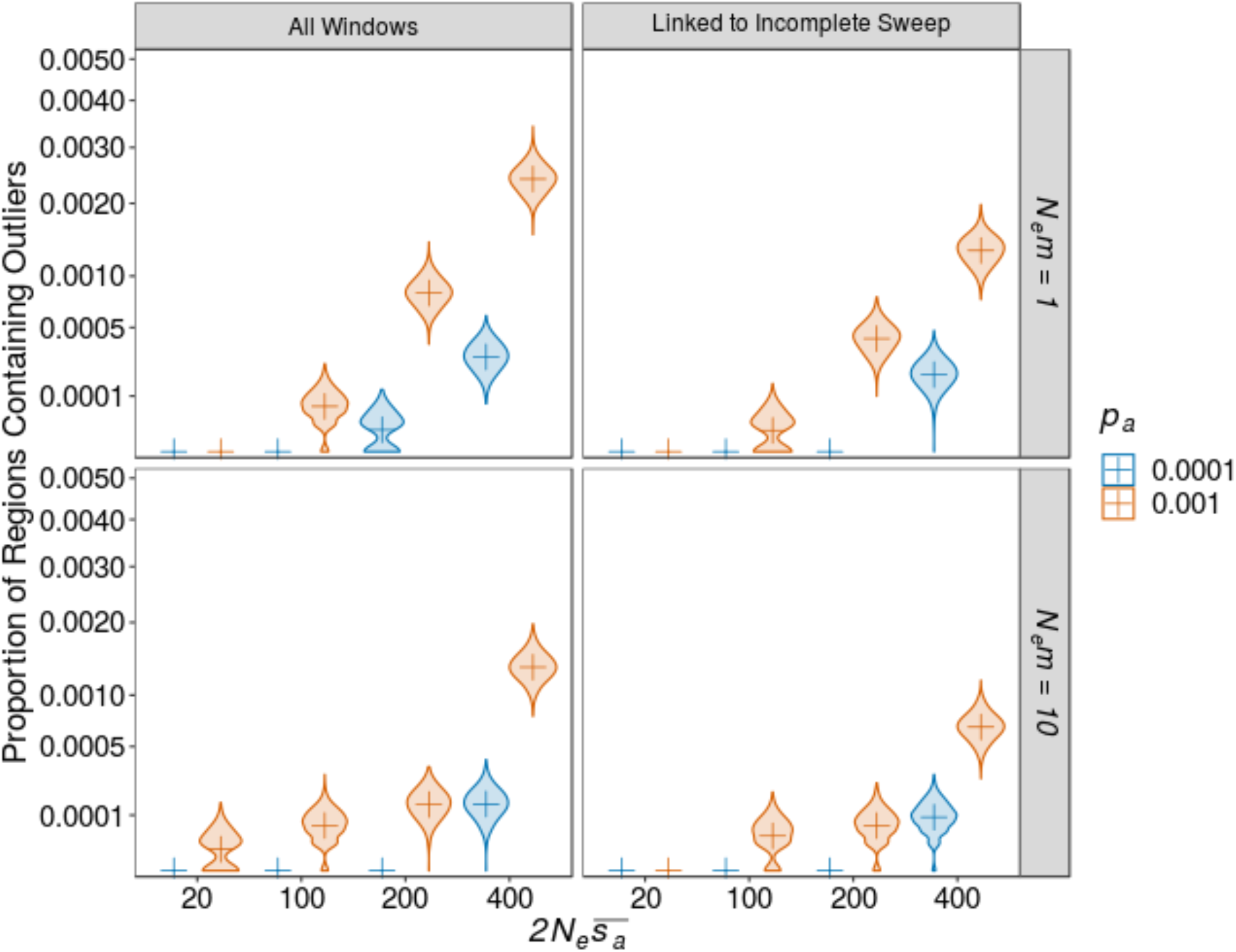
The proportion of analysis windows that contain *F_ST_* outliers in parapatric populations. Allele-frequency weighted *F_ST_* was calculated for 10,000 bp analysis windows centred on simulated “gene-like” regions. Plusses indicate the point estimate, and violins indicate the distribution of 1,000 bootstraps samples from 2,000 simulation replicates.

**Figure S3.**
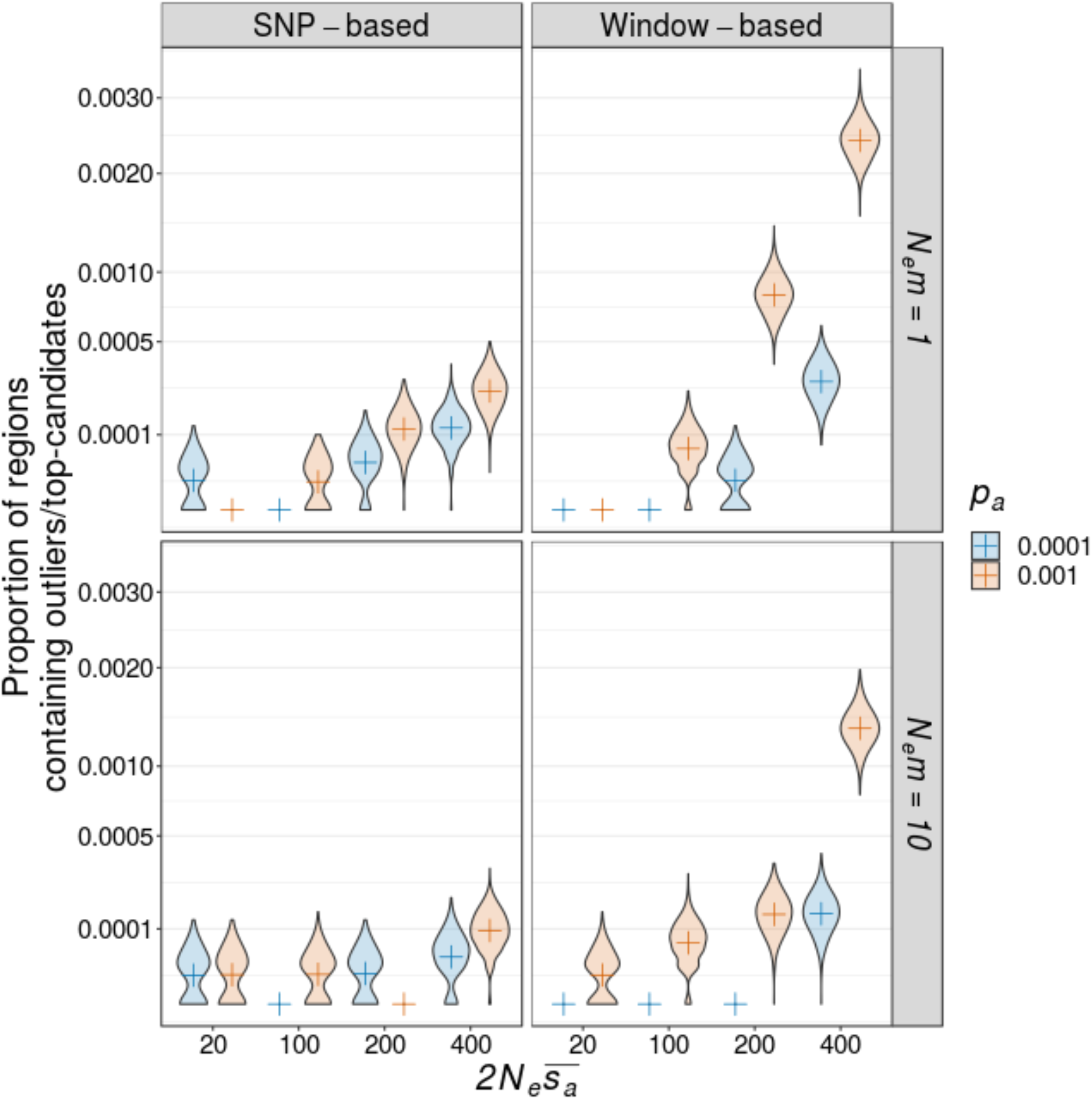
A comparison of SNP-based and analysis window-based detection methods. Top-candidates are regions with a significant excess of high *F_ST_* SNPs. Window-based data are plotted in Figure 3. Plusses indicate the point estimate and violins indicate the distribution of 1,000 bootstraps samples from 2,000 simulation replicates.

**Figure S4.**
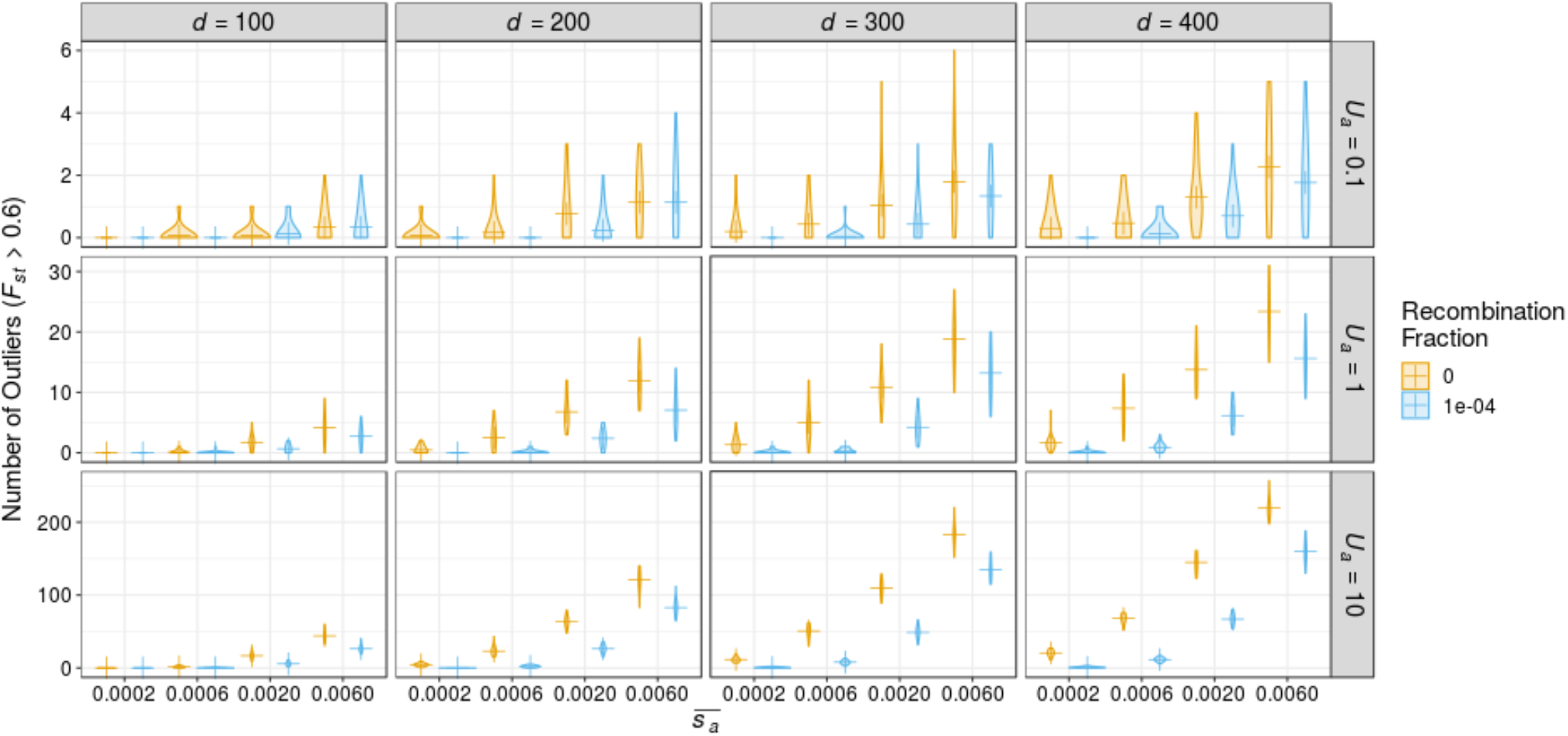
The number of outlier SNPs in the stepping-stone model. The plusses indicate the mean number of outliers. Outliers are those with *F_ST_* greater than the 99.999th percentile of the distribution from neutral simulations. *U_a_* is the number of new advantageous mutations which occur each generation, *d* is the distance, in number of demes, separating the focal demes and 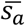 is the mean of an exponential distribution of advantageous mutational effects.

## Appendix 1

### The proportion of a sweep’s duration in which it may induce differentiation between two points in a one-dimensional range

Consider the spread of advantageous alleles in a spatially continuous, one-dimensional environment. The classical results of Fisher (1937) and Kolmogorov et al., (1937) showed that advantageous alleles spread through space in wave-like fashion at a constant speed of 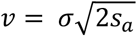, assuming a Gaussian dispersal kernel with standard deviation *σ*. Similar results have been reported for a variety of dispersal kernel (Ralph & Coop, 2010). Here, we derive a simple approximation for the proportion of the fixation time in which differentiation may be induced between two populations in a one-dimensional range.

Consider a species dispersed with uniform density over a 1-dimensional landscape of length R. Populations *A* and *B* are located at positions *x*(*A*) and *x*(*B*), respectively and are separated by a distance of *Δx*. Here, we derive the proportion of a sweep’s sojourn time in which it may induce genetic differentiation between regions *A* and *B*. We assume that advantageous mutations spread in wave-like fashion, and that *Δx* is much greater than the length of the wave front. Furthermore, we assume that differentiation may be induced if the wave is somewhere between regions *A* and *B*.

For any advantageous mutation occurring on the landscape that sweeps to fixation, there are two possibilities; that it occurs between *A* and *B*, which happens with probability 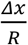, or that the mutation occurs either side of the interval between *A* and *B*, this happens with a probability of 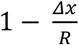. If the beneficial mutation occurs somewhere between *A* and *B*, on average the sweep has to travel a distance of 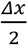 before it reaches both populations. In this case, the time it takes to reach both populations is thus 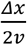. If the mutation lands outside of the region between *A* and *B*, the allele has to travel a distance of *Δx*, and this takes a time of 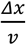. The mean time an advantageous mutation may generate differentiation is thus,

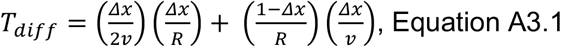

which reduces to,

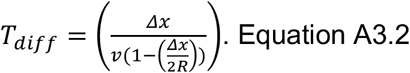

The mean time for a complete sweep depends on the length of the species’ range. For any new mutation introduced at a random point in a species’ range, the average distance from the point of origin to the farthest range limit is 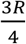. The expected fixation time is thus approximately 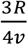. The proportion of a sweep’s fixation time (*P_diff_*), in which it may induce differentiation between *A* and *B* is

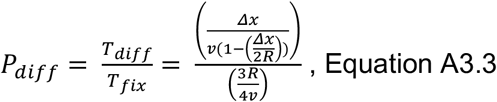

which simplifies to

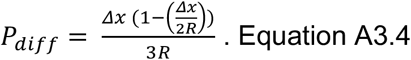

Wave speed cancels out in Equation A3.4 because the wave is moving at a constant speed.

## Appendix 2

### The number of ongoing selective sweeps in a panmictic populations

In natural populations, the true underlying population structure is difficult to ascertain. As a result, it would be difficult to develop an analytical model for the number of ongoing sweeps in a structured population. Instead, we derive the number of alleles sweeping in a single panmictic population.

Consider a Wright-Fisher population of *N* diploids subject to recurrent mutations. Our model assumes free recombination and as such we do not explicitly model deleterious variants. New mutations occur at rate *μ* per base-pair per generation. For *η_a_* sites in the genome, a proportion of new mutations generate beneficial alleles (*p_a_*). These alleles are advantageous with selective effects drawn from an exponential distribution with mean 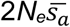. We use the fixation time of an advantageous allele from (Ewens, 1979):

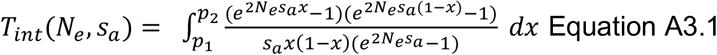

To obtain an estimate of the expected time a sweeping allele spends at intermediate frequency during its sojourn, we integrate from *p*_1_ to *p*_2_. Given a distribution of fitness effects for new mutations, there will be a distribution of expected times to fixation. To obtain the average of the distribution of times to fixation, we integrate over the DFE, giving:

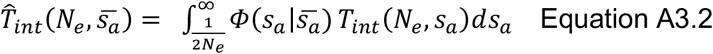

where 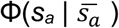 is the probability density function of the distribution of fitness effects. The number of new advantageous mutations which arise each generation per individual, *U_a_*, is a product of the mutational target size *η* and mutation rate for advantageous alleles *μ_a_* (*μ_a_* = *μp_a_*).

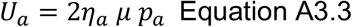

The selection coefficients and corresponding fixation probabilities for the new advantageous mutations will vary according to a distribution of fitness effects. Integrating over the DFE, we obtain the expected number of new advantageous alleles, destined for fixation, which arise each generation (*V_a_*),

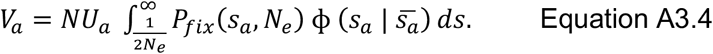

In this study we assume that the DFE for advantageous mutations is an exponential distribution with mean 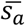.

The expected number of ongoing sweeps, *X_a_* can be calculated as:

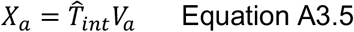

### The proportion of substitutions attributable to positive selection, *α*

If all adaptive substitutions are due to selection acting on *de novo* mutations, the proportion of substitutions attributable to positive selection (*α*) can be obtained by integrating over the relevant portions of the distribution of fitness effects (DFE:

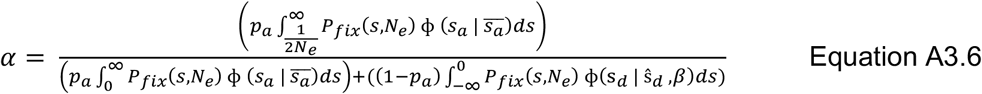

Where *P_fix_*(*s*) is the probability of fixation for a new mutation with selection coefficient *s*. *P_fix_* is calculated using Kimura’s formula:

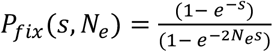

In the case of advantageous mutations with 2*N_e_s_a_* > 1, *P_fix_* is approximately *2s*. The integral in the numerator is from 1/2*N_e_* to ∞ because weakly advantageous mutations with fitness effects *2N_e_s_a_* ≥ 1 have fixation probabilities similar to neutral ones and thus it cannot be determined whether substitutions of such alleles are due to positive selection or drift.

A Mathematica workbook containing an implementation of this model is available at https://github.com/TBooker/GlobalAdaptation.

